# Cortical activation associated with motor preparation can be used to predict the freely chosen effector of an upcoming movement and reflects response time: An fMRI decoding study

**DOI:** 10.1101/295345

**Authors:** Satoshi Hirose, Isao Nambu, Eiichi Naito

## Abstract

Motor action is prepared in the human brain for rapid initiation at the appropriate time. Recent non-invasive decoding techniques have shown that brain activity for action preparation represents various parameters of an upcoming action. In the present study, we demonstrated that a freely chosen effector can be predicted from brain activity measured using functional magnetic resonance imaging (fMRI) before initiation of the action. Furthermore, the activity was related to response time (RT). We measured brain activity with fMRI while 12 participants performed a finger-tapping task using either the left or right hand, which was freely chosen by them. Using fMRI decoding, we identified brain regions in which activity during the preparatory period could predict the hand used for the upcoming action. We subsequently evaluated the relationship between brain activity and the RT of the upcoming action to determine whether correct decoding was associated with short RT. We observed that activity in the supplementary motor area, dorsal premotor cortex, and primary motor cortex measured before action execution predicted the hand used to perform the action with significantly above-chance accuracy (approximately 70%). Furthermore, in most participants, the RT was shorter in trials for which the used hand was correctly predicted. The present study showed that preparatory activity in cortical motor areas represents information about the effector used for an upcoming action, and that well-formed motor representations in these regions are associated with reduced response times.

**Highlights:**

- Brain activity measured by fMRI was used to predict freely chosen effectors.
- M1/PMd and SMA activity predicted the effector hand prior to action initiation.
- Response time was shorter in trials in which effector hand was correctly predicted.
- Freely chosen action is represented in the M1/PMd and SMA.
- Well-formed preparatory motor representations lead to reduced response time.

## 1. Introduction

Prior preparation can facilitate subsequent motor actions. Indeed, response times (RTs) are known to decrease when specific instructions are provided prior to the initiation of a required action (Rosenbaum, 1980; Jahanshahi et al., 1992). These findings suggest that “motor preparation” before the initiation of an action can improve reaction times.

Previous studies have investigated the neural mechanisms underlying motor preparation by examining changes in neural activity (preparatory activity) between instruction presentation and action initiation (preparatory period) in the brains of both humans and non-human primates (Shenoy et al., 2013; Turella and Lingnau, 2014; DiRusso et al., 2017). Such studies have revealed that preparatory activity reflects the parameters of an upcoming movement. For example, studies on non-human primates have demonstrated that the parameters of an upcoming movement (e.g., movement direction) are represented in cortical motor areas, such as the primary motor cortex (M1; Evarts and Tanji, 1976; Tanji and Evarts, 1976), dorsal premotor cortex (PMd; Godschalk et al., 1981), and supplementary motor area (SMA; Tanji et al., 1980).

Additional primate studies have revealed that neural activity associated with movement parameters may influence preparedness to execute an upcoming movement, thereby affecting RT (see Riehle and Requin, 1993; Bastian et al., 2003).

Human preparatory activity was originally identified using electroencephalography (EEG; Kornhuber and Deecke, 2016; Shibasaki et al., 1980; Libet et al., 1983; Shibasaki and Hallett, 2006). As in the non-human primate brain, it has been suggested that human preparatory activity also represents information regarding movement parameters (Coles et al., 1988; Vaughan et al., 1968; Eimer, 1998; Haggerd and Eimer, 1999). However, because of limitations in the spatial and temporal resolution of non-invasive measurement techniques, the precise associations between brain activation and specific action parameters, as well as the regions involved, remain to be fully elucidated.

Recently, functional magnetic resonance imaging (fMRI) decoding has made significant contributions to identifying parameters of upcoming actions represented in human preparatory activity (Bode and Haynes, 2009; Gallivan et al., 2011a; Gallivan et al., 2011b; Gallivan et al., 2013; Nambu et al., 2015). In fMRI decoding, the contents of mental processes are predicted based on brain activity, using machine learning methods (Haynes and Rees, 2006; Norman et al., 2006; Formisano et al., 2008; Pereira et al., 2009). Recent fMRI studies have successfully identified specific components of upcoming actions such as grasp shape (Gallivan et al., 2011a), reaching direction (Gallivan et al., 2011b), the effector used for the upcoming action (Gallivan et al., 2013), and patterns of sequential finger movements (Nambu et al., 2015) in the parietal and frontal regions, which include the SMA, M1, and PMd. Furthermore, a recent study demonstrated that freely chosen (internally motivated) grasp shapes can be predicted based on preparatory activity in multiple frontal and parietal brain regions (Ariani et al., 2015).

To further investigate the neural representations of upcoming actions in preparatory activity, we determined whether and where in the brain a freely chosen effector (the left or right hand) is represented during a preparatory period. Furthermore, we investigated whether action representations in specific regions are associated with the preparedness of the action, as indicated by improvement in RT.

In the present study, we used fMRI to measure the brain activity of 12 participants during a finger-tapping task in which the effector (left or right hand) was freely chosen by the participants. We used fMRI decoding analyses to identify brain regions in which the preparatory activity measured prior to action initiation could be used to predict the freely chosen effector. We first performed decoding analysis using all voxels in the brain (whole-brain decoding) to determine whether information about the freely chosen effector existed in the preparatory activation measured with fMRI. We then evaluated the weight distribution of the decoding model to identify brain regions that primarily represented the information. Following the whole-brain decoding, *post hoc* decoding analyses were performed using voxels in the identified brain regions (regional decoding) to determine whether the effector hand can be predicted based only on their activity.

Finally, we evaluated the relationship between decoding accuracy and RT, hypothesizing that RTs would be relatively shorter for correctly predicted responses than for incorrectly predicted responses. This hypothesis was based on previous studies involving monkeys (Churchland et al., 2006) and mice (Hasegawa et al., 2017), which demonstrated that better preparedness (i.e., shorter RT) is associated with specific patterns of preparatory neural activity, and that the pattern of neural activity is more variable when the action is poorly prepared.

## 2. Methods

### 2.1. Participants

Twelve (8 men and 4 women) healthy, right-handed (laterality quotient > 80; Oldfield, 1971) participants (age [mean ± standard deviation]: 25.6 ± 4.4 years) were recruited in the present study. The study was approved by the Ethical Committee of the National Institute of Information and Communications Technology and conducted in accordance with the Declaration of Helsinki (1975), and all participants provided written informed consent prior to the experiment.

### 2.2. Task procedure

The participants rested comfortably in the supine position in an fMRI scanner, with their arms oriented parallel to the torso and forearms pronated and supported by a cushion, allowing them to relax their arms. During fMRI scanning, the participants were instructed to move their fingers only, without moving their wrists.

Visual stimuli were projected onto a screen in the scanner such that the participants could view them in a mirror positioned in front of their eyes. Throughout the experiment, a fixation cross was presented in the center of the screen, and the participants were asked to maintain their gaze on this point to avoid unnecessary eye movements.

Three black circles were presented for 4 s at the beginning of each trial (Decision Period; Figure 1). We pseudo-randomly selected these three circles from four options, each of which was assigned to indicate the left middle, left index, right middle, or right index finger. During the Decision Period, participants were asked to freely choose one of the three circles and to prepare to move the corresponding finger. After the Decision Period, the circles disappeared. Subsequently, after a variable delay (Delay Period; 0, 2 or 4 s), the fixation cross turned red (go signal), and the participants tapped with the selected finger on an MR-compatible button box (Current Design Inc., Philadelphia, PA) for 4 s (Execution Period). The participants were instructed to initiate the movement as quickly as possible upon the presentation of the go signal and to tap the button as many times as possible during the 4-s Execution Period. The button box was placed just below the fingers, allowing the participants to easily press the button without moving their wrists.

**Figure 1:**
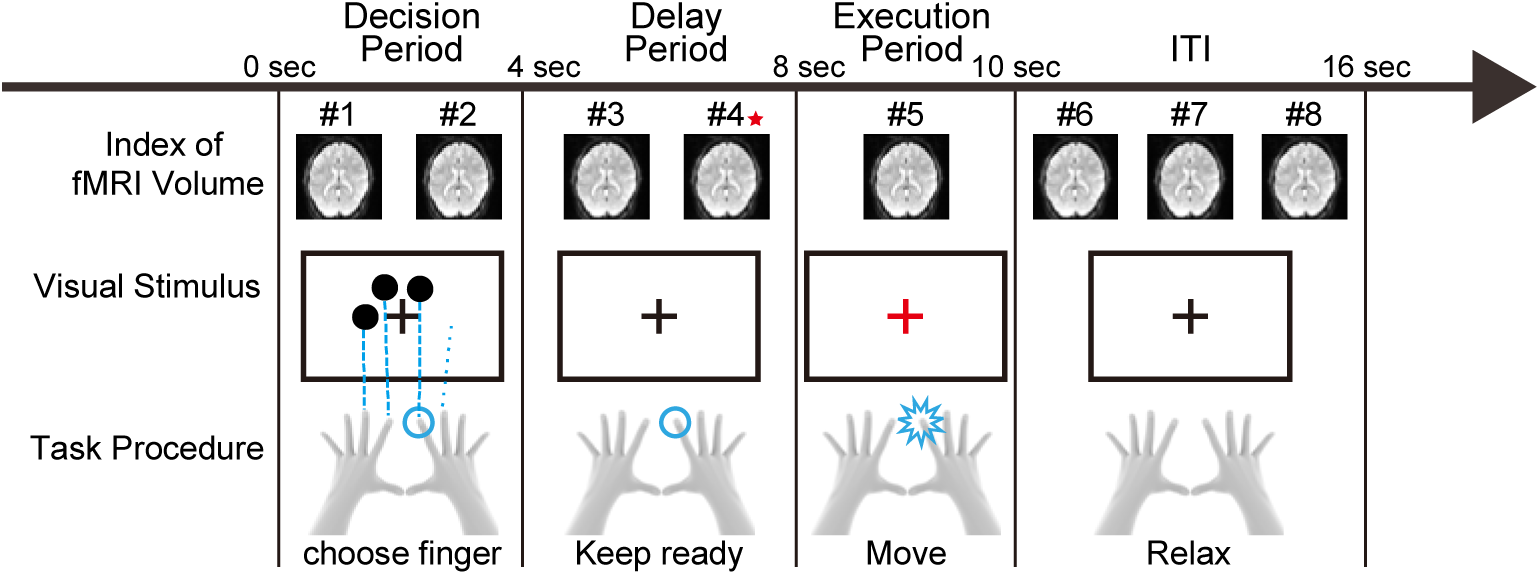
Task procedure for a target trial. Each trial began with a 4-s presentation of three black circles, which were randomly chosen from among four candidates whose positions corresponded to the right index, right middle, left index, and left middle fingers (Decision Period). During this period, participants freely selected one of the three indicated fingers and prepared to move the selected finger (cyan circle). After the Decision Period, the circles disappeared. Subsequently, after a 4-s delay (Delay Period), the fixation cross turned red, and the participants tapped with the selected finger as many times as possible for 4 s (Execution Period). After executing the movement, the participants relaxed for the inter-trial interval (ITI). We analyzed eight volumes, acquired in the Decision Period (Volumes #1 and #2), Delay Period (Volumes #3 and #4), Execution Period (Volume #5) and ITI (Volumes #6, #7, and #8).

Each session consisted of 24 trials, comprising 15 target trials with a 4-s delay, 3 catch trials with a 2-s delay, and 6 catch trials without a Delay Period. In the 24 trials in a session, each of the four possible sets of three circles during the Decision Period appeared in 6 trials. The order of the circles and the duration of the Delay Period was randomized independently. Trials were separated by an inter-trial interval (ITI) of 6 s. Each session also included a 16-s rest period before the first trial (pre-resting period), and another 10-s rest period after the last trial (post-resting period). In total, we collected 187 functional images in each session, including pre-resting and post-resting periods (repetition time [TR] = 2,000 ms). Each participant completed a total of 150 target trials: 15 target trials for each of the 10 sessions.

### 2.3. fMRI measurement

We used a 3.0-T SIEMENS scanner (Trio Tim) with a 12-channel head coil to obtain T1-weighted anatomical images (MP-RAGE; TR: 2,250 ms; echo time [TE]: 3.06 ms; inversion time [TI]: 900 ms; flip angle: 9 degree, field-of-view [FOV]: 256 × 256 mm; voxel size: 1.0 mm × 1.0 mm × 1.0 mm) and functional T2*-weighted echo-planar images (EPIs; 64 × 64 matrix; pixel size: 3.0 mm × 3.0 mm; TE: 30 ms; slice thickness: 4 mm; gap between slices: 1 mm; number of slices: 30; voxel size: 3.0 mm × 3.0 mm × 5.0 mm; FOV: 192 mm × 192 mm × 150 mm [whole brain within FOV]. We collected 187 functional volumes in each session.

### 2.4. fMRI data preprocessing and trial selection

We excluded the first five functional volumes in each session from the analysis to allow for magnetization equilibrium, following which the remaining images were realigned to correct for head movement and co-registered to each participant’s anatomical image.

To minimize differences in the magnitude of the fMRI voxel values across sessions, we calculated the percentage increase in the fMRI signal in each voxel from its mean value in each session. The percent increase was calculated by dividing each voxel value in an fMRI volume by its mean value across the 182 volumes of the session. Finally, a temporal high-pass filter (Butterworth filter) with a cutoff frequency of 1/128 Hz was applied to the data obtained in each session to remove low-frequency drift.

fMRI data were preprocessed using the MATLAB “Multi Voxel Pattern Classification (MVPC) toolbox” (Hirose et al., 2015; http://www2.nict.go.jp/bnc/hirose/mvpc/index.html), which uses functions of Statistical Parametric Mapping Software (SPM; http://www.fil.ion.ucl.ac.uk/spm/).

Decoding analyses were performed for brain activity in the 150 target trials (catch trials were excluded from these analyses). We also excluded target trials in which no button-press occurred, as well as those in which participants moved two or more fingers during the Execution Period, since we could not distinguish which finger movement had been prepared. Eleven target trials were excluded in one participant, and five in another. One target trial each was excluded in two additional participants, while no trials were excluded in the remaining seven participants.

### 2.5. General methodology for decoding analyses

In the decoding analyses described below (Sections 2.6 and 2.8), a “decoder” was trained using a supervised classification algorithm, with a training dataset composed of pairs of multi-voxel patterns (e.g., percentage increases for the voxels) and their corresponding labels (e.g., hand used for the upcoming movement). In the present study, we focused on hand selection. Thus, trials in which the participants selected the right index or the right middle finger were assigned the same label (label = 1), while trials in which participants selected either of the left fingers were assigned a label of −1. The goal of the decoding analysis was to generate a decoder that could accurately predict the label (i.e., hand) from the multi-voxel patterns of fMRI signals that were not included in the training dataset (test data).

To evaluate the prediction accuracy of the decoder (decoding accuracy), we performed a leave-one-session-out cross-validation. Specifically, we trained the classifiers using the trials from nine sessions (training dataset) and used the trials of the remaining session to evaluate prediction accuracy (test dataset). This validation was performed for all 10 possible session combinations (i.e., 10-fold cross validation). Prior to classifier training in each validation test, we normalized the voxel values by first linearly transforming the values in the training dataset into z-scores, with a mean of 0 and standard deviation of 1. The mean and standard deviation (SD) of each voxel in the training dataset were used in the same way to transform the voxel values in the test dataset. We defined the mean prediction accuracy across the 10 validation tests as the decoding accuracy for each participant.

We used an iterative sparse logistic regression (iSLR; Hirose et al., 2015) algorithm for pattern classification. The iSLR is designed based on Bayesian sparse logistic regression (SLR; Yamashita et al., 2008). In iSLR, multiple SLR classifiers are trained with a training dataset, and the final label prediction is obtained by combining the predictions of these SLR classifiers. The number of SLR classifiers is a free parameter that controls the sparseness of the model (i.e., how many voxels are selected). In the present study, the value of this parameter was determined by performing leave-one-session-out (9-fold) cross validation within the training dataset (nested cross validation). The search space included natural numbers between 1 and 100. The means and standard deviations of the parameters and the number of voxels selected by iSLR are included in the Supplementary Table.

For optimization, we used the iSLR algorithm implemented in the MVPC toolbox (Hirose et al., 2015; http://www2.nict.go.jp/cinet/bnc/hirose/mvpc/index.html), which uses functions from the Sparse Logistic Regression Toolbox (Yamashita et al., 2008; SLR Toolbox; http://www.cns.atr.jp/~oyamashi/SLR_WEB.html).

A significance threshold of *p* = 0.05 was used for all statistical analyses. Unless otherwise specified, the statistical significance was evaluated using two-tailed permutation tests with Bonferroni correction for multiple comparisons (i.e., the *p* values were multiplied by the number of comparisons). In the present study, we report these *p* values with the correction factors in the subscript (e.g., “*p*_8_”).

In the present study, we mainly used the permutation test, which is theoretically preferable to the t-test for fMRI decoding studies, as the t-test requires the data to be normally distributed, which is not always satisfied in fMRI decoding studies (Nichols and Holmes, 2001; Golland and Fischl, 2003; Etzel et al., 2008; Smith and Muckli, 2010; Chen et al., 2011; Galliivan et al., 2011a,b). The methodological details of the permutation tests are described in Appendix A.

### 2.6. Whole-brain decoding

We first conducted the decoding analysis using all voxels within the entire brain. The brain voxels were identified using the following procedure, which is implemented in SPM. First, the mean value of all fMRI signals measured during the experiment (10 sessions) was calculated for each voxel (signal intensity). Then, the average of the signal intensities across all voxels was calculated (global mean). Finally, voxels whose signal intensity was greater than 80% of the global mean were identified as brain voxels, including white matter, cerebrospinal fluid, gray matter, and subcortical nuclei. The number of brain voxels was 31,957 ± 1,785 (mean ± SD across participants).

We conducted the decoding analyses separately for each of eight fMRI volumes: two volumes from the Decision Period (#1 and #2 in Figure 1), two from the Delay Period (#3 and #4), one from the Execution Period (#5), and three from the ITI (#6, #7, and #8).

One-sample t-tests were used to compare the mean decoding accuracy across participants against chance (50%) for each of the eight volumes. The *p* values were corrected for multiple comparisons using Bonferroni’s method (i.e., multiplied by 8; significance threshold: *p*_8_ < 0.05). To verify that movement artifacts did not influence the results, we also performed the same decoding analysis using voxels outside the brain (Appendix B).

We further assessed statistical significance using permutation tests (Appendix A1). Due to their high computational cost, permutation tests were performed for volume #4 only, as this volume was most likely to represent preparatory brain activity (Section 3.1 in Results) and did not contain motion artifacts (Appendix B).

### 2.7. Identifying relevant brain regions

To identify the preparatory representation of the hand that would be used in the upcoming movement, we analyzed the spatial location of the voxels selected by iSLR (Hirose et al., 2015) for each individual participant. We then evaluated the consistency of our findings across participants.

For individual analyses, we identified the voxels that were selected in more than half of the 10 validation tests (= selected voxels). The locations of these voxels were transformed into Montreal Neurological Institute (MNI) coordinates, and an image with 2 mm isotropic voxels representing the selected voxels was created. We then smoothed each participant’s image by adding the voxels that surrounded each selected voxel (≤ 4 mm between centers) as “smoothed selected voxels.” This was done to compensate for slight differences across participants in the spatial locations of the selected voxels, which may have been caused by errors in spatial normalization or individual neuroanatomical differences.

To examine the spatial consistency of the selected voxels across participants, we counted the number of participants (NP score) in whom each voxel was a smoothed selected voxel. Voxels with NP scores > 2 were defined as “relevant voxels”. Finally, clusters of more than 94 contiguous relevant voxels were defined as “relevant clusters”. This threshold for the cluster size was determined using the permutation test as described in Appendix A.4. The SPM Anatomy toolbox (Eickhoff et al., 2007; http://www.fz-juelich.de/inm/inm-1/DE/Forschung/_docs/SPMAnatomyToolbox/SPMAnatomyToolbox_node.html) was used for anatomical identification of the relevant clusters.

### 2.8. Decoding analysis from the identified brain regions (regional decoding)

From the whole-brain decoding, we identified three relevant clusters (see Section 3.2 in Results, Figure 2B). Two of these clusters were located in the lateral parts of the precentral gyri in both hemispheres, appearing to correspond to the bilateral M1 and PMd. The remaining cluster covered the medial parts of the bilateral precentral gyri and most likely corresponded to the SMA. Although voxels in these regions were used for effector prediction, such results do not definitively indicate that these brain regions included information regarding the selected effector, as these regions contained only a subset of the selected voxels. Theoretically, a subset of selected voxels need not always contain relevant information (Haufe et al., 2014). Thus, to verify that the fMRI data from each identified region (M1/PMd and SMA) contained relevant information, we performed *post hoc* decoding analyses using only voxels within the relevant clusters.

**Figure 2:**
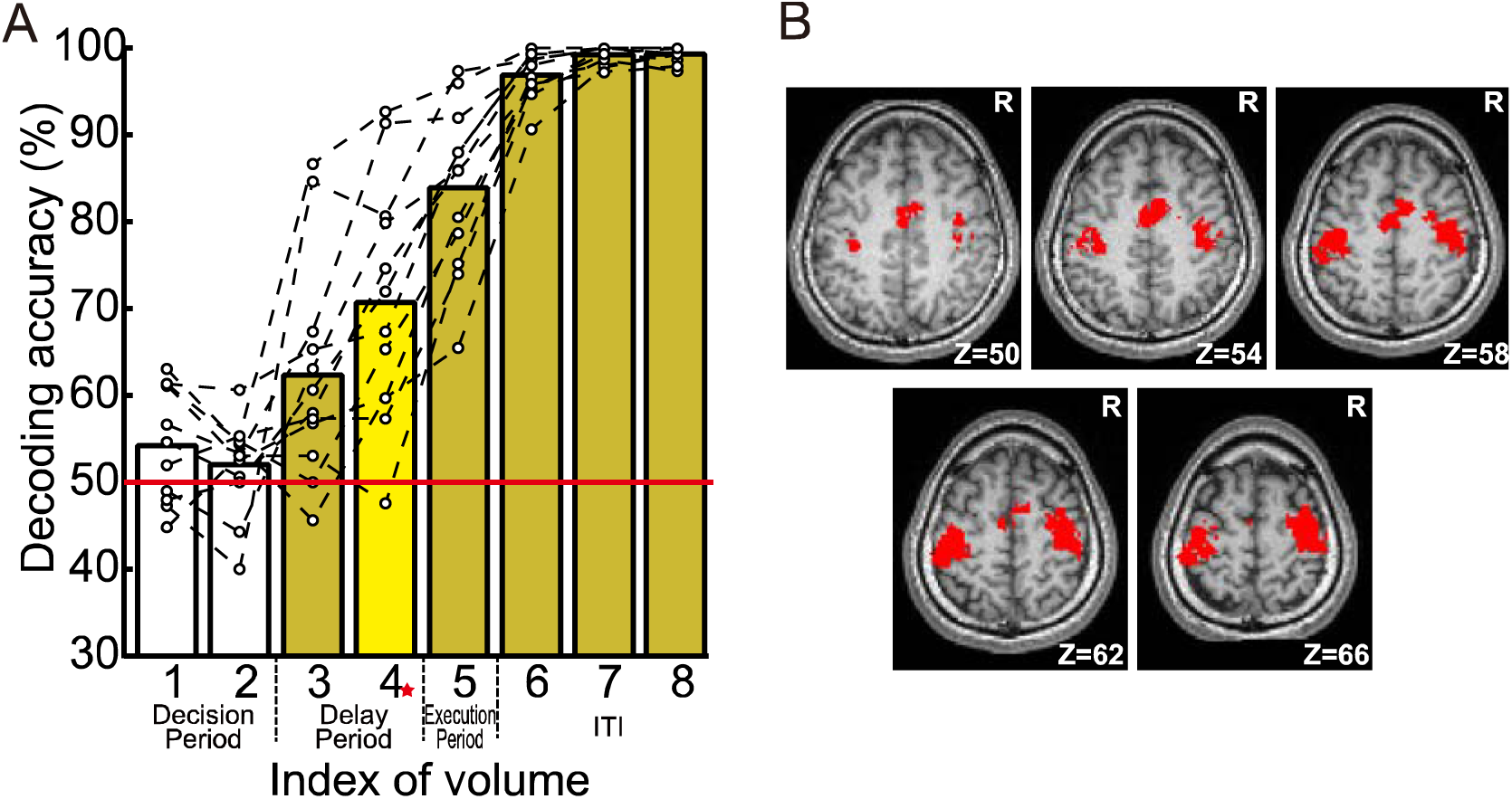
Results of the whole-brain decoding. A) Mean (bars) and each participant’s (white dots connected with a dashed line) decoding accuracies. The dark yellow bars indicate the volumes for which t-tests revealed significantly higher-than-chance mean decoding accuracy (50%; red line), while the light yellow bar indicates the volume for which a permutation test (conducted only for this volume) revealed significantly higher-than-chance mean decoding accuracy. White bars indicate lack of significance in the t-test. B) Relevant clusters identified in group analysis for Volume #4 (red-starred in A). Red sections indicate the relevant clusters, which were defined as clusters of relevant voxels whose sizes were above threshold. The map is overlaid on the normalized anatomical image of a participant.

First, we defined two region-of-interest (ROI) images representing the lateral and medial frontal brain regions (M1/PMd ROI, SMA ROI) in the MNI standard brain. The ROIs included all voxels in the relevant clusters and certain surrounding voxels (distance ≤ 4 mm between centers). The surrounding voxels were included to minimize the possibility of excluding relevant voxels. The ROI images were then transformed into each participant’s individual brain coordinates. Finally, using all voxels in each ROI, we evaluated the decoding accuracies using the same procedures as described for the whole-brain decoding (M1/PMd and SMA decoding; see Section 2.6). If the predictions remained accurate even when we only used voxels in each ROI, the ROI was considered likely to contain relevant effector information. To examine this possibility, we compared the decoding accuracies obtained from the M1/PMd and SMA decoding with chance (50%) using the permutation test described in Appendix A.1.

In addition, we performed the same decoding analysis using the brain voxels outside of the ROIs only (OUT-OF-ROI decoding; Pessoa and Padmala, 2006; Etzel et al., 2013). In this case, we expected that the decoding accuracy would decrease relative to that observed for the whole-brain decoding. To evaluate this prediction, we compared the decoding accuracy obtained from OUT-OF-ROI decoding with that obtained from the whole-brain decoding (Appendix A.2).

### 2.9. Relationship between RT and decoding correctness

Finally, we hypothesized that the RT, which was defined as time from the Go signal (when the fixation cross turned red) to the first response of the finger, would be shorter in trials where a decoder made correct predictions (correctly-decoded trials) than in those where the decoder made incorrect predictions (incorrectly-decoded trials). This hypothesis was evaluated for the M1/PMd, SMA and OUT-OF-ROI decoding. First, we calculated the median RT across trials for each participant, separately for the correctly- and incorrectly-decoded trials. The differences between the trial groups were then evaluated using the permutation test (Appendix A.3). As stated in the Introduction, we expected that if preparatory activity in these regions reflects preparedness for the action, RT should be shorter for correctly-decoded trials than for incorrectly decoded trials.

## 3. Results

### 3.1. Whole-brain decoding

Figure 2A shows the decoding accuracies obtained from the whole-brain decoding. The mean decoding accuracies (± SD) during the Decision Period were 54.3 ± 6.4% for Volume #1 and 52.0 ± 5.4% for Volume #2. These were both near chance (50%). In contrast, the decoding accuracy increased during the Delay Period (62.4 ± 12.5% for Volume #3 and 70.7 ± 13.8% for Volume #4). The decoding accuracy increased further during the Execution Period and during the ITI (83.9 ± 9.5% for Volume #5, 96.9 ± 2.7% for Volume #6, 99.2 ± 1.0% for Volume #7, and 99.3 ± 0.9% for Volume #8).

T-tests showed significantly above-chance decoding accuracy in Volumes #3 through #8 (#3: t_11_ = 3.4, *p*_8_ = 0.04; #4: t_11_ = 5.2, *p*_8_ = 0.0024; #5: t_11_ = 12.4, *p*_8_ = 6.6 × 10^−7^; #6: t_11_ = 60.5, *p*_8_ = 2.5 × 10^−11^; #7: t_11_ = 168, *p*_8_ = 3.3 × 10^−16^; #8: t_11_ = 193, *p*_8_ = 7.2 × 10^−23^). In contrast, the decoding accuracy was not significantly greater than chance during the Decision Period (#1: t_11_ = 2.33, *p*_8_ = 0.32; #2: t_11_ = 1.29, *p*_8_ > 1). The non-parametric permutation test also confirmed the statistical significance of above-chance decoding accuracy for Volume #4 (*p* < 10^−6^; Appendix A.1).

The better-than-chance decoding accuracies during the Delay Period (Volumes #3 and #4) are most likely driven by preparatory brain activity, because the activity during this period was temporally separated from contamination by motion artifacts and movement-related brain activity. Indeed, we can suppose, based on the shape of the hemodynamic response (Friston et al., 1998), that the measured responses in Volume #3 likely reflected the early component of the blood-oxygen-level-dependent (BOLD) hemodynamic response caused by neural activity during the Decision Period. The activity in Volume #4 corresponds well to the peak of the BOLD response.

When we analyzed the voxels outside the brain (see Appendix B for details), the decoding accuracy was nearly at chance for Volume #4. On the other hand, the decoding accuracy significantly exceeded chance (*p*_8_ < 0.05) in Volumes #5 (Execution Period) and #6 (first volume during ITI), in which the button press occurred in almost all trials. These results indicate that noise components associated with finger movements (e.g., head movement artifacts) affected the fMRI signals even in voxels outside the brain during the Execution Period (Volume #5) and the beginning of ITI (Volume #6), but not during the second half of the Decision Period (Volume #4).

### 3.2 Identifying brain regions representing essential information

We identified three clusters of relevant voxels (one-tailed *p*_48_ < 0.05; Figure 2B). Two of these clusters were located in the lateral parts of the precentral gyri in both hemispheres (right: 934 voxels, (x, y, z) = (35 mm, –20 mm, 64 mm), left: 834 voxels, (x, y, z) = (–39 mm, –27 mm, 63 mm)) and the remaining cluster covered the medial parts of the bilateral precentral gyri (cluster size = 439 voxels, MNI coordinates of the center of gravity: (x, y, z) = (1 mm, –8 mm, 56 mm)). The lateral clusters corresponded to M1 and the PMd, while the medial cluster likely corresponded to the SMA. Other identified clusters did not reach significance (≤ 142 voxels; Appendix Figure A.4).

### 3.3 Regional decoding

In the regional decoding (Figure 3), the mean decoding accuracies were significantly better than chance (*p*_*2*_ < 10^−5^; Appendix A.1) in both the M1/PMd decoding (73.0 ± 12.8%) and the SMA decoding (67.7 ± 10.6%). Significantly lower decoding accuracy was observed in the OUT-OF-ROI decoding than was achieved for the whole-brain decoding (*p* = 0.0039; Appendix A.2).

**Figure 3:**
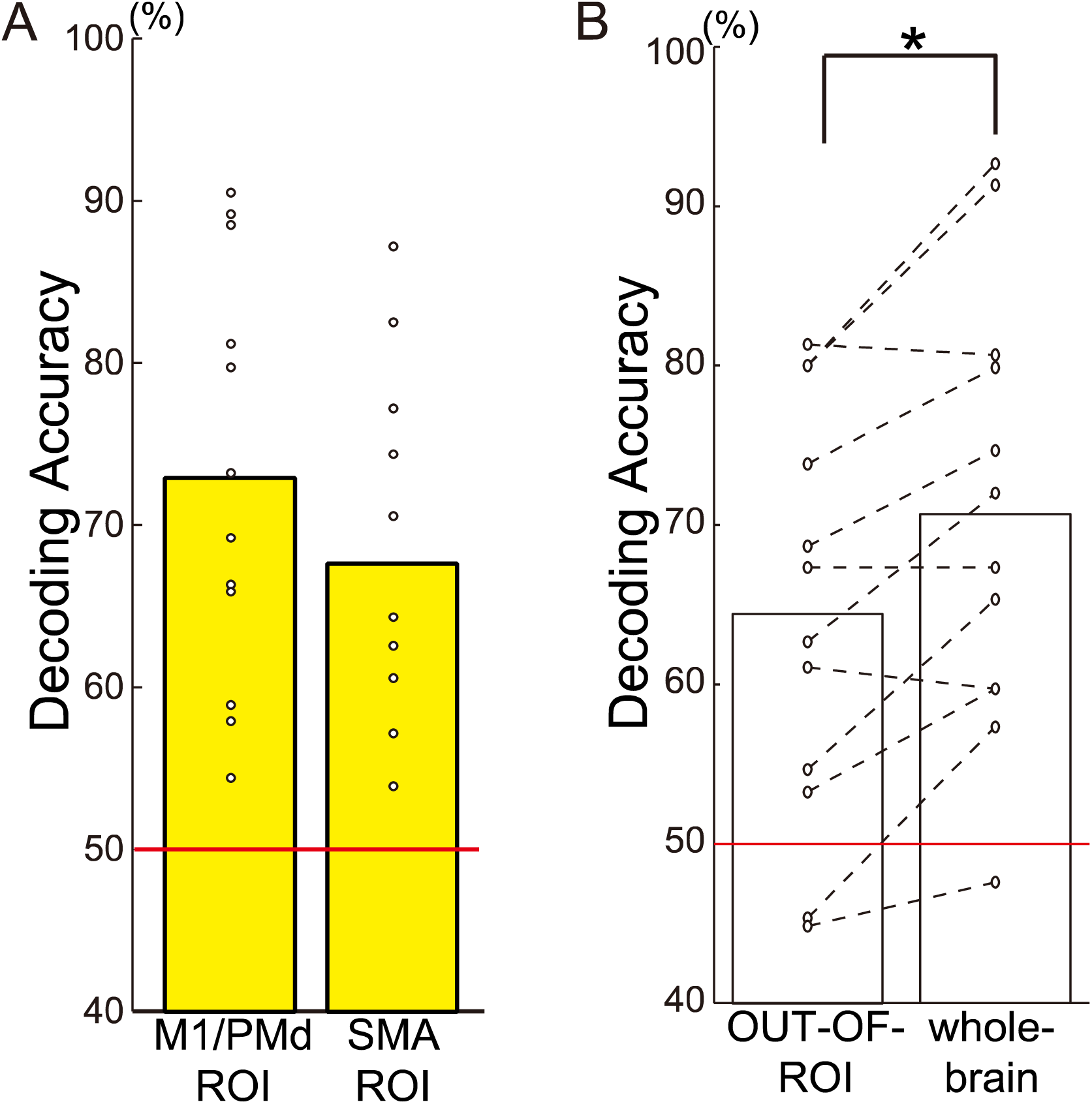
Results of regional decoding. A: Decoding accuracies obtained from the M1/PMd and SMA decoding. Yellow bars indicate that the mean decoding accuracy was significantly higher than the level of chance (50 %; red line). B) Decoding accuracy obtained from the OUT-OF-ROI decoding, in which voxels outside of the M1/PMd and SMA ROIs were used for decoding. Results from the whole-brain decoding are shown again for comparison. The asterisk indicates the significant difference between the whole-brain and OUT-OF-ROI analyses (*p* < 0.05). Structure abbreviations: M1, Primary motor cortex; PMd, dorsal premotor cortex; SMA, supplementary motor area.

The greater-than-chance decoding accuracies obtained from the M1/PMd and SMA decoding verified the presence of effector information in each of these areas, while the significantly lower decoding accuracy obtained from OUT-OF-ROI decoding further indicated that the M1/PMd and/or SMA ROIs contained primary information for predicting the effector.

The regional decoding was a *post hoc* analysis of the whole-brain decoding based on the premise that the results of the whole-brain decoding were correct. Consequently, the analyses inherently involved a problem of circularity or “double dipping” (Kriegeskorte et al., 2009). However, we were able to replicate the results using an analysis with a non-circular ROI selection procedure with no concerns about double dipping, that is, a procedure in which the ROIs for each participant was determined on the basis of results from the remaining 11 participants (results not shown).

### 3.4 Relationship between RT and decoding correctness

Finally (Figure 4), we observed that the RT tended to be longer for incorrectly-decoded trials than for correctly-decoded trials, for both the M1/PMd (Figure 4A; correct: 466 ± 100 ms, incorrect: 489 ±102 ms; mean difference: 23 ms) and SMA decoding (Figure 4B; correct: 467 ± 100 ms, incorrect: 481 ± 103 ms; mean difference: 14 ms). However, no such finding was observed in the OUT-OF-ROI decoding (Figure 4C; correct: 471 ± 101 ms, incorrect: 472 ± 104 ms; mean difference: –1.5 ms). Indeed, we observed a significant difference in mean RT between correctly- and incorrectly-decoded trials for the M1/PMd decoding (*p*_*3*_ = 0.013) and a near-significant trend for the SMA decoding (*p* = 0.039; uncorrected for multiple comparisons), but no significant differences were observed for the OUT-OF-ROI decoding (*p*_*3*_ > 1) (Appendix A.3).

**Figure 4:**
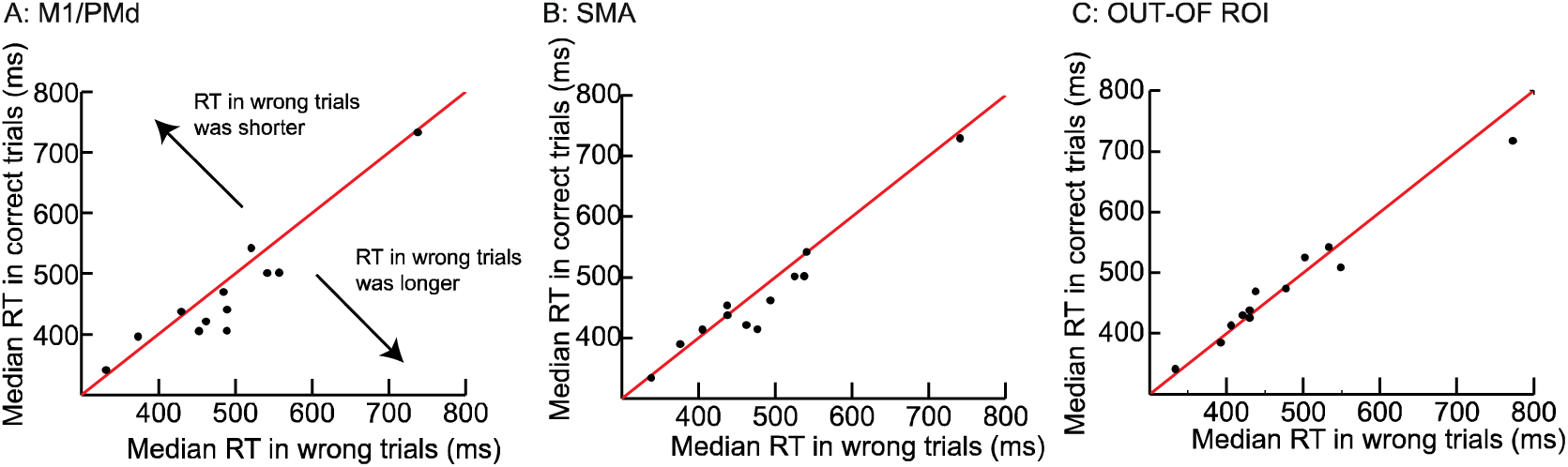
Median response times (RTs) in the correctly- and incorrectly-decoded trials. Median RT in trials for which the correct predictions were made by the decoder (correctly-decoded trials; vertical axes) was plotted against that for incorrectly-decoded trials (horizontal axis). Panels A, B, and C represent results from the M1/PMd, SMA, and OUT-OF-ROI decoding, respectively. Each dot represents results from one participant. The red line indicates the case in which the median RTs are identical between the correctly- and incorrectly-decoded trials, while the distance from the red line indicates the difference in RT between correct and incorrectly-decoded trials. Dots lower than the red line represent participants whose RT in incorrectly-decoded trials was longer than that in correctly-decoded trials. Abbreviations: M1, primary motor cortex; PMd, dorsal premotor cortex; SMA, supplementary motor area; ROI, region of interest.

## 4. Discussion

In the present study, we demonstrated that we could predict the freely chosen motor effector used in a delayed finger-tapping task from fMRI signals measured before action execution (Figure 2A). Such predictions could be achieved based on preparatory activities in cortical motor areas (M1/PMd and SMA; Figure 2B), which therefore contain information regarding the choice of the effector for an upcoming action prior to its execution (Figure 3). Decoding using voxels outside of the brain (skull decoding; Appendix B) further supported the conclusion that the successful decoding was unaffected by artifacts associated with finger movements. Moreover, analyses of the sequences of hand choices confirmed that the successful prediction of the effector hand could not be explained by behavioral choice biases (Supplementary Material 1). Finally, we also showed that the RT becomes shorter in correctly-decoded trials than in incorrectly-decoded trials (Figure 4).

### 4.1. Freely-chosen effector information is represented in M1/PMd and SMA

The whole-brain decoding identified relevant clusters in M1/PMd and SMA (Figure 2B), and the *post hoc* regional decoding for the M1/PMd and SMA further validated the conclusion that the information used for the effector prediction was represented in these cortical motor areas (Figure 3A). The decoding accuracy became significantly lower when we excluded the voxels in these ROIs (OUT-OF-ROI; Figure 3B). These results clearly indicate that the activity in the cortical motor areas (M1/PMd and SMA) represented primary information to allow prediction of the freely chosen motor effector used in an upcoming action.

Previous fMRI decoding studies reported that action parameters can be predicted from PMd, M1 or SMA (shape of grasp: Gallivan et al., 2011a, reaching direction: Gallivan et al., 2011b, patterns of sequential finger movements: Nambu et al., 2015). Specifically, an effector (left or right hand) specified by an external cue (forced-choice) can be decoded from preparatory activity in these regions (Gallivan et al., 2013). Our present results generally support the notion that preparatory activity in human M1, PMd and SMA represents parameters of an upcoming motion, by adding evidence that these cortical motor areas may also represent information about a freely-chosen effector.

In a gross perspective, there is a dominance of the contralateral hemisphere in the preparatory activity for an action. Studies that measured event-related potentials (ERP) using EEG (Barrett et al. 1986; Cui and Deecke 1999; McAdam and Seales 1969) have suggested that the cerebral hemisphere contralateral to a motor effector (e.g. hand finger) is more activated during motor preparation. Furthermore, a study that measured electrocorticographic (ECoG) signals from one patient (Ikeda et al., 1992) and an fMRI study (Michelon et al., 2006) both reported that preparatory activity in the contralateral M1 and SMA is modestly higher, although the ipsilateral corresponding regions also increase their activities. Thus, the successful effector prediction in our study (Figures 2, 3A) was most probably achieved because the decoder emphasized the differences in activation between the two hemispheres by giving appropriate weights to the voxels in each hemisphere. Indeed, in both the M1/PMd and SMA ROIs, the decoder gave positive weights mainly to voxels in the left hemisphere, and negative weights mainly to voxels located in the right hemisphere (Supplementary Material 2 and Supplementary Figure). Since higher activity of positively weighted voxels pushes the decoder toward effector prediction of the right hand, while higher activity of negatively weighted voxels pushes the decoder toward effector prediction of the left hand, the results suggest contralateral dominance in the preparatory activity of M1/PMd and SMA when the brain freely chooses a motor effector.

Ariani et al. (2015) reported that three different types of movements of the right hand (precision grip, power grip or reaching without hand movement) could be predicted from the preparatory activation of the contralateral M1, PMd and SMA but not from the ipsilateral regions. Together with our present findings, this suggests that contralaterally-dominant activation likely reflects a contralaterally-dominant computational process in the preparation of an upcoming hand action.

### 4.2. Relation between RT and brain activity

We found that RT became shorter in trials where the hand that was used was correctly predicted from the cortical motor activity (Figure 4). To the best of our knowledge, this is the first study to demonstrate that fMRI decoding can be used to reveal a relationship between preparatory motor representations and length of RT. Since this prediction was achieved based on multi-voxel fMRI patterns, the result indicates that such patterns should differ between correctly-decoded and incorrectly-decoded trials. Further studies are needed to determine the precise neuronal mechanisms that determine rapid initiation of action. However, we raise the following two possibilities.

First, we raise the possibility that the amplitude of the BOLD signal is greater in trial with shorter RTs. This view seems to be in accordance with the rise-to-threshold model (Hanes and Schall, 1996; Erlhagen and Schöner, 2002). In this model, the preparatory activity of a group of neurons involved in an upcoming action increases until it reaches the threshold for generating the action. When the threshold is crossed, the action is initiated. Thus, the RT may be shorter in a trial where the preparatory activity level is closer to the threshold, which may also allow for easier decoding because of the higher signal strength.

Second, we may point out the possibility that the variability in neural activity is lower in trials with shorter RTs. This idea accords with the suggestion that variability of neuronal activity for well-prepared movement is relatively lower, while that for poorly-prepared movement is greater (Churchland et al., 2006; Shenoy et al., 2011; Hasegawa et al., 2017). If this is the case, decoding would be less accurate in poorly-prepared trials, even when mean neuronal activity is the same. This view seems to be supported by a recent transcranial magnetic stimulation (TMS) study (Klein-Flügge et al., 2013), which showed that variance in M1 excitability prior to the onset of a finger movement is lower in trials with shorter RTs.

## 5. Summary and Conclusions

One concern regarding the current study is that the limited number of participants (12) may restrict the generalizability of the results. However, our study demonstrated that the fMRI signal measured before action execution can be used to predict the effector hand freely chosen for the upcoming movement, and that this effector information is primarily represented in the M1/PMd and SMA during the preparatory phase of action execution. Furthermore, our findings demonstrate that the neural representations in these brain regions are directly related to the motor preparation that enables the rapid initiation of the chosen action. Further studies are required to determine how M1/PMd and SMA receive signals associated with internal decision-making from upper stream brain regions, such as pre-SMA, anterior cingulate cortices, dorsolateral prefrontal cortex (Rae et al., 2014; Soon et al., 2008), and use them to form neural representations essential for the rapid initiation of a chosen action.

## Appendix A Permutation test

Recent studies have suggested that the permutation test is preferable to the t-test in fMRI decoding studies, as the t-test requires the data to be normally distributed, which is not always satisfied in such studies (Nichols and Holmes 2001; Golland and Fischl, 2003; Etzel et al., 2008; Smith and Muckli, 2010; Chen et al., 2011; Galliivan et al 2011a,b). Thus, in the present study, we mainly used permutation tests to determine statistical significance. However, permutation tests were performed only for Volume #4 during the whole-brain decoding, because of the high computational cost of this process.

We permuted the attribution of the data (e.g., left or right hand for each trial) so that the permuted result was equally likely to be observed under the null hypothesis (Nichols and Holmes 2001). Repeated permutation allows for the empirical estimation of the distribution under the null hypothesis (null distribution) of a value (e.g., mean decoding accuracy), which is the target of the statistical test.

The specific methods and the results of the permutation tests for each statistical test are described below.

### A.1. Permutation test for comparison of decoding accuracy with chance (50%)

This test was used for the decoding accuracy obtained from the whole-brain decoding for Volume #4 and for decoding accuracies obtained from M1/PMd and SMA decoding.

#### Method

We set the null hypothesis as, *“The preparatory brain activation was unrelated to the label (left or right hand)”* and permuted the labels of the trials. The labels (left or right hand) in each session were randomly permuted for each participant. Using this permuted dataset, we then performed the decoding analysis and calculated the decoding accuracy via leave-one-session-out cross validation, using the same procedure as described in the Methods section (Section 2.6). This was repeated 100 times for each participant, to obtain 100 decoding accuracies under the null hypothesis.

Finally, we randomly selected one of 100 decoding accuracies for each participant and then calculated the mean decoding accuracy across participants. This random selection was repeated 1,000,000 times to obtain 1,000,000 mean decoding accuracies under the null hypothesis. The statistical significance of the true decoding accuracy, which was obtained from the analysis without permutation, was tested using this distribution.

#### Results

Appendix Figure A.1 shows the estimated probability density of mean decoding accuracy obtained from the whole-brain decoding for Volume #4 and for decoding accuracies obtained from the M1/PMd and SMA decoding (black bars). The true mean decoding accuracies without permutation were 70.7 %, 73.0 % and 67.7% for the whole-brain, M1/PMd and SMA decoding respectively. Among 1,000,000 repetitions, no permutation result was above the true decoding accuracy for any analyses (uncorrected one-tailed *p* < 10^−6^). Thus, we may conclude that these decoding accuracies were significantly higher than chance (two-tailed *p* < 10^−5^ for the whole-brain decoding; corrected two-tailed *p*_2_ < 10^−5^ for regional decoding).

**Appendix Figure A.1:**
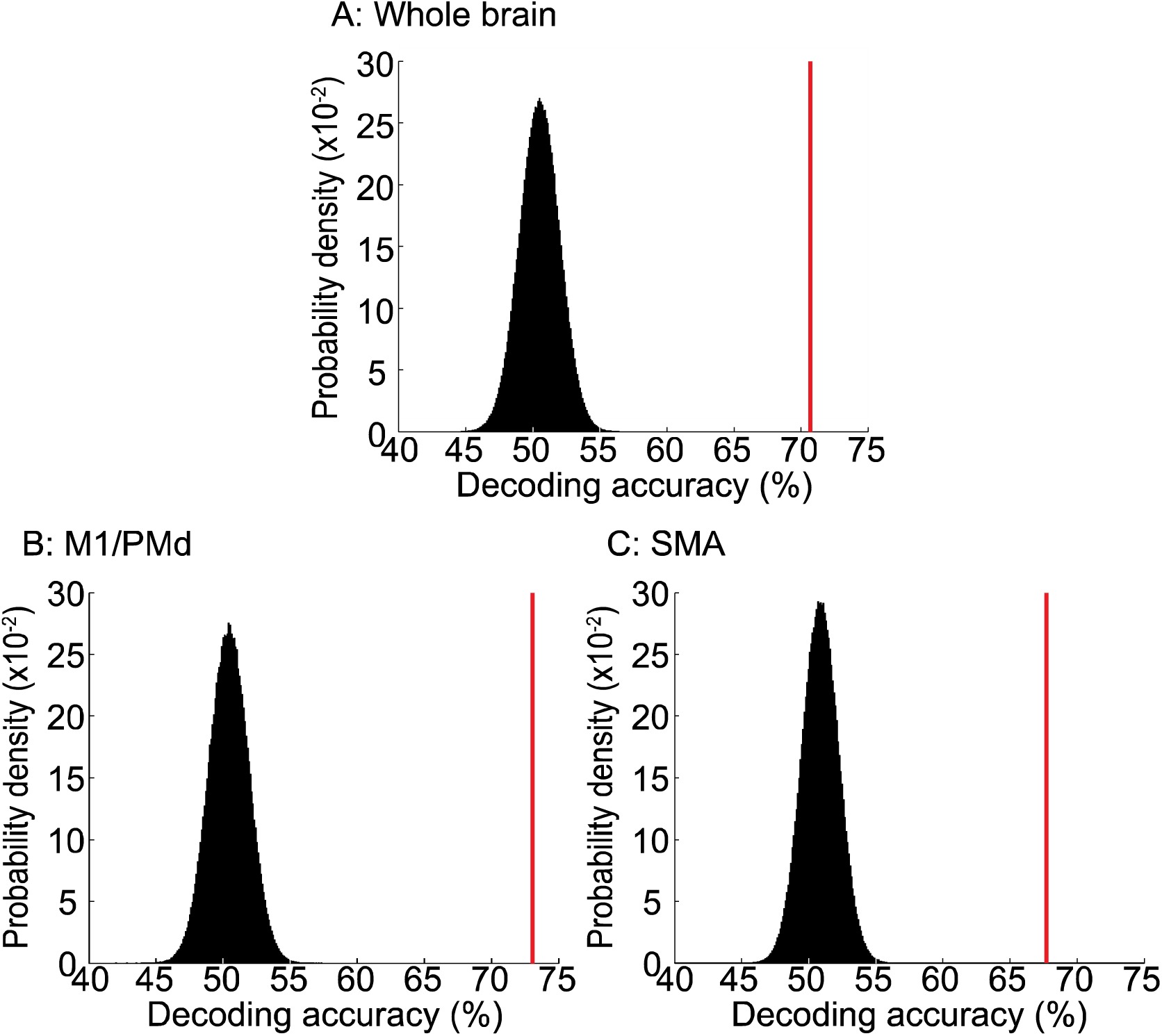
Estimated probability density of mean decoding accuracies under the null hypothesis, *“The preparatory brain activation was unrelated to the label (left or right hand).”* The panels represent results from (A) the whole-brain, (B) the M1/PMd, or (C) the SMA decoding. Black bars indicate the estimated probability density for each bin (bin size = 0.1 %). Red vertical bars indicate the true decoding accuracies without permutation (A: 70.7%, B: 73.0%, C: 67.7%).

### A.2. Permutation test for comparison of decoding accuracies

This test was used to evaluate the difference in decoding accuracies between the whole-brain and the OUT-OF-ROI decoding.

#### Method

We set the null-hypothesis as *“The decoding accuracies are identical for the whole-brain and the OUT-OF-ROI decoding”* and permuted the attribution of decoding accuracy. Namely, decoding accuracy obtained from the whole-brain decoding for each participant was attributed as decoding accuracy for OUT-OF-ROI, and vice versa. Using all 4096 (2^12^) combinations, we achieved 4096 values for the mean difference of the decoding accuracy under the null hypothesis. The statistical significance of the true difference in decoding accuracy, which was obtained from the analysis without permutation, was tested using this distribution.

#### Results

Appendix Figure A.2 shows the estimated probability density of the mean difference in the decoding accuracies between the whole-brain and the OUT-OF-ROI decoding under the null hypothesis (black bars) and the true difference without permutation (6.3%). In 8 of 4096 repetitions, the difference was larger than or equal to the true difference (uncorrected one-tailed *p* = 0.002). Thus, we may conclude that the decoding accuracy was significantly higher for the whole-brain than the OUT-OF-ROI decoding (two-tailed *p* = 0.0039).

**Appendix Figure A.2:**
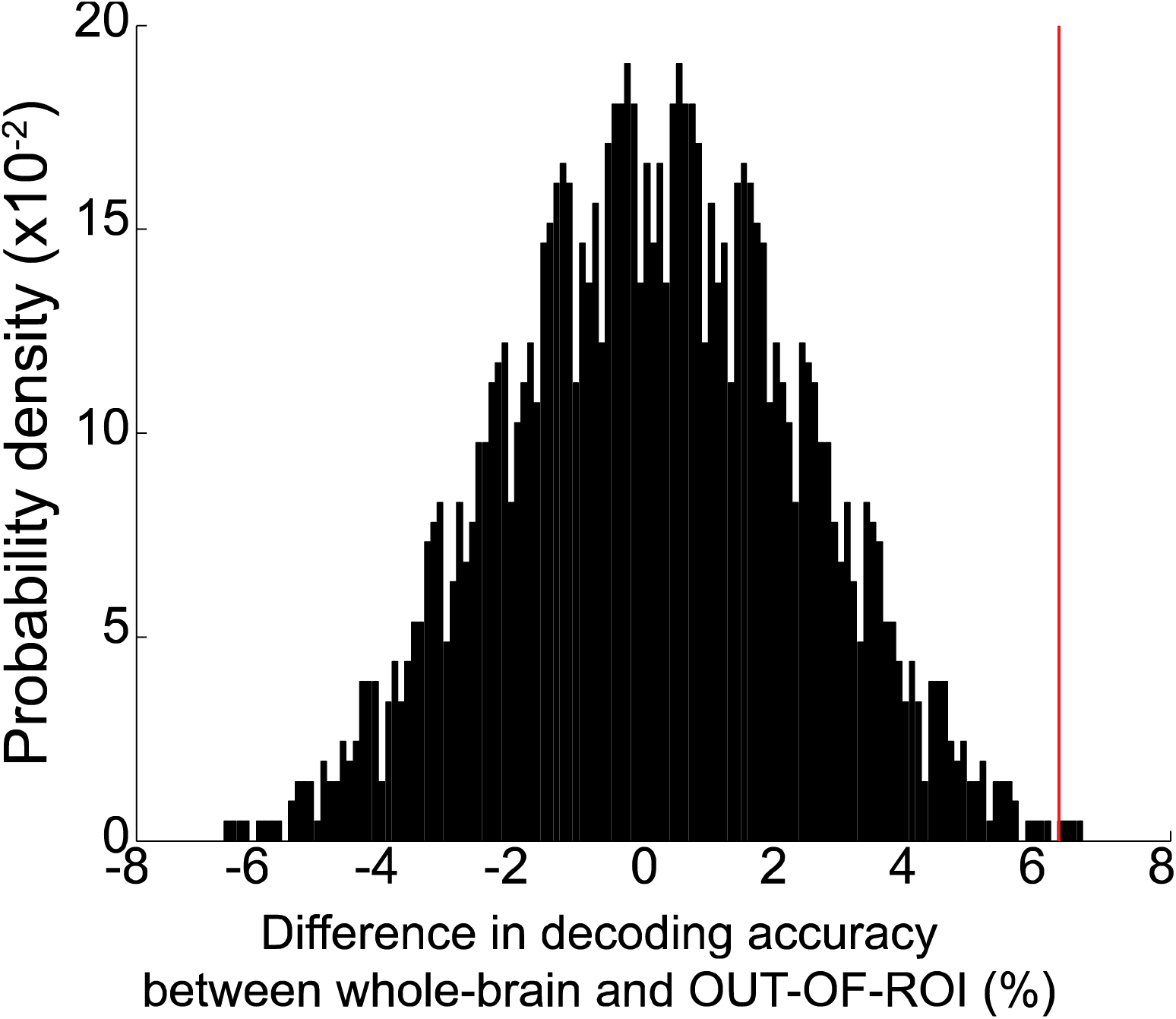
Estimated probability density of the mean difference in the decoding accuracies between the whole-brain decoding and the OUT-OF-ROI decoding, under the null hypothesis, *“The decoding accuracies are identical for the whole-brain and OUT-OF-ROI decoding.”* Positive values on the horizontal axis indicate that the mean decoding accuracy was higher for the whole-brain decoding than for OUT-OF-ROI decoding. Black bars indicate the probability density for each bin (bin-size = 0.1 %). The red vertical bar indicates the true difference without permutation (6.3%).

### A.3. Permutation test for comparison of RTs

This test was used to evaluate differences in RT between correctly- and incorrectly-decoded trials.

#### Method

We set the null-hypothesis as *“The correctness of decoding is unrelated to the RT”* and permuted the correctness (correct or incorrectly-decoded trial) of the trials. Specifically, we randomly permuted this attribution for each participant and computed the mean (across participants) difference of the median RT using the procedure described in the Methods section (Section 2.9). This was repeated 1,000,000 times, and the statistical significance of the mean decoding accuracy was tested using this distribution.

#### Results

Appendix Figure A.3 shows the estimated probability density of the mean RT difference between correct and incorrectly-decoded trials (black bars) and the true mean RT without permutation (M1/PMd: 23 ms, SMA: 15 ms, OUT-OF-ROI: –1.5 ms). In 0.21, 1.97, and 57.7% of the permutations, respectively, the RT difference was larger than the true difference (uncorrected one-tailed *p* = 0.0021, *p* = 0.0197, *p* = 0.423). Thus, the significance of the difference was confirmed for the M1/PMd (corrected two-tailed *p*_*3*_ = 0.013), while values approaching significance were noted for the SMA (two-tailed *p* = 0.039 without Bonferroni correction). No significant differences were noted for the OUT-OF-ROI (corrected two-tailed *p*_*3*_ > 1).

**Appendix Figure A.3:**
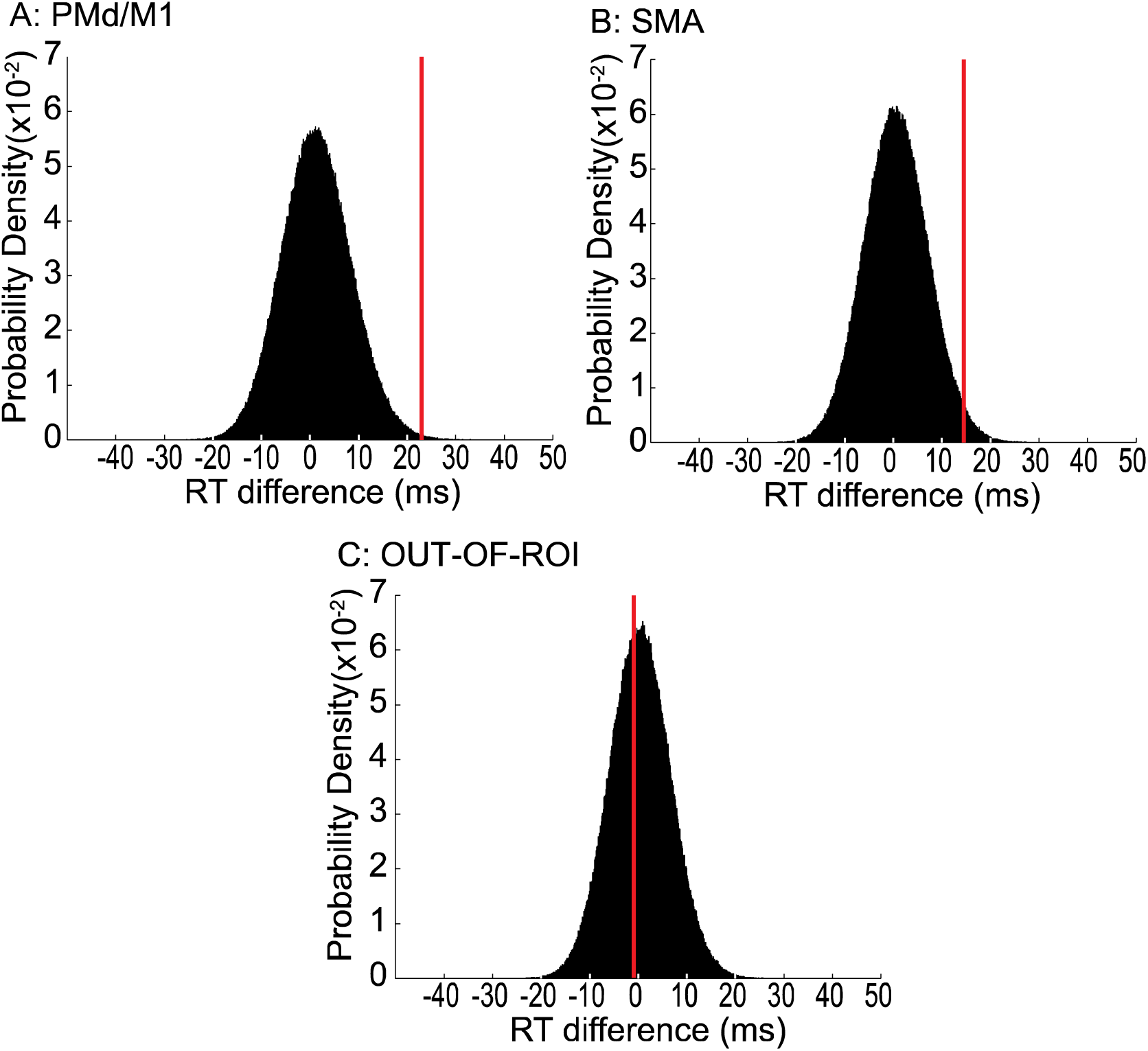
Estimated probability density of the mean RT difference between correctly- and incorrectly-decoded trials under the null hypothesis, *“The correctness of decoding is unrelated to the RT.”* Each panel represents results obtained from one ROI (A: M1/PMd, B: SMA, C: OUT-OF-ROI). Positive values on the horizontal axis indicate that the mean RT was larger in the incorrectly-decoded trials than in the correctly-decoded trials. Black bars indicate the probability density for each bin (bin size = 0.1 ms). Red vertical bar indicates the true mean RT without permutation (A: 23 ms, B: 15 ms, C: –1.2 ms).

### A.4: Permutation test for cluster size

This test was used to evaluate the size of the clusters of the relevant voxels

#### Method

We set the null-hypothesis as *“The selected voxels were located randomly in the brain.”* Recall that “the selected voxels” indicate the voxels that were selected in more than half of the 10 validation tests for each participant. In this test, we randomly permuted the locations of the relevant voxels. For each participant, we repeated the random permutation 100 times and prepared 100 images representing the randomly permuted selected voxels. These images were normalized and smoothed using the procedure described in the Methods section (Section 2.7).

Then, we randomly selected one image from among the 100 images of each participant and identified the relevant voxels whose NP score was > 2 (see Section 2.7) to calculate the maximum cluster size. The random selection was repeated 1,000,000 times to obtain 1,000,000 maximum sizes of the clusters under the null hypothesis. When the size of a true cluster obtained from the analysis without permutation was larger than the upper 5% point of the null distribution, we regarded the cluster as significant (a relevant cluster).

#### Result

Appendix A.4 shows the estimated cumulative probability density of maximum cluster size (black bars) and a histogram of the sizes of the true clusters without permutation (red bars). Because we identified 48 clusters of relevant voxels from the analysis without permutation, we set the threshold to p_48_ = 0.05 (uncorrected p = 0.001), which yielded a value between 142 and 143 (vertical dotted line). Thus, clusters whose sizes were > 142 were regarded as significant (relevant clusters). From the analysis without permutation, three relevant clusters (i.e. size > 142) were identified. Two of these relevant clusters were located in the lateral parts of the precentral gyri in both hemispheres (right and left M1/PMd in Appendix Figure A.4). The remaining cluster covered the medial parts of the bilateral precentral gyri (SMA in Appendix Figure A.4).

**Appendix Figure A.4:**
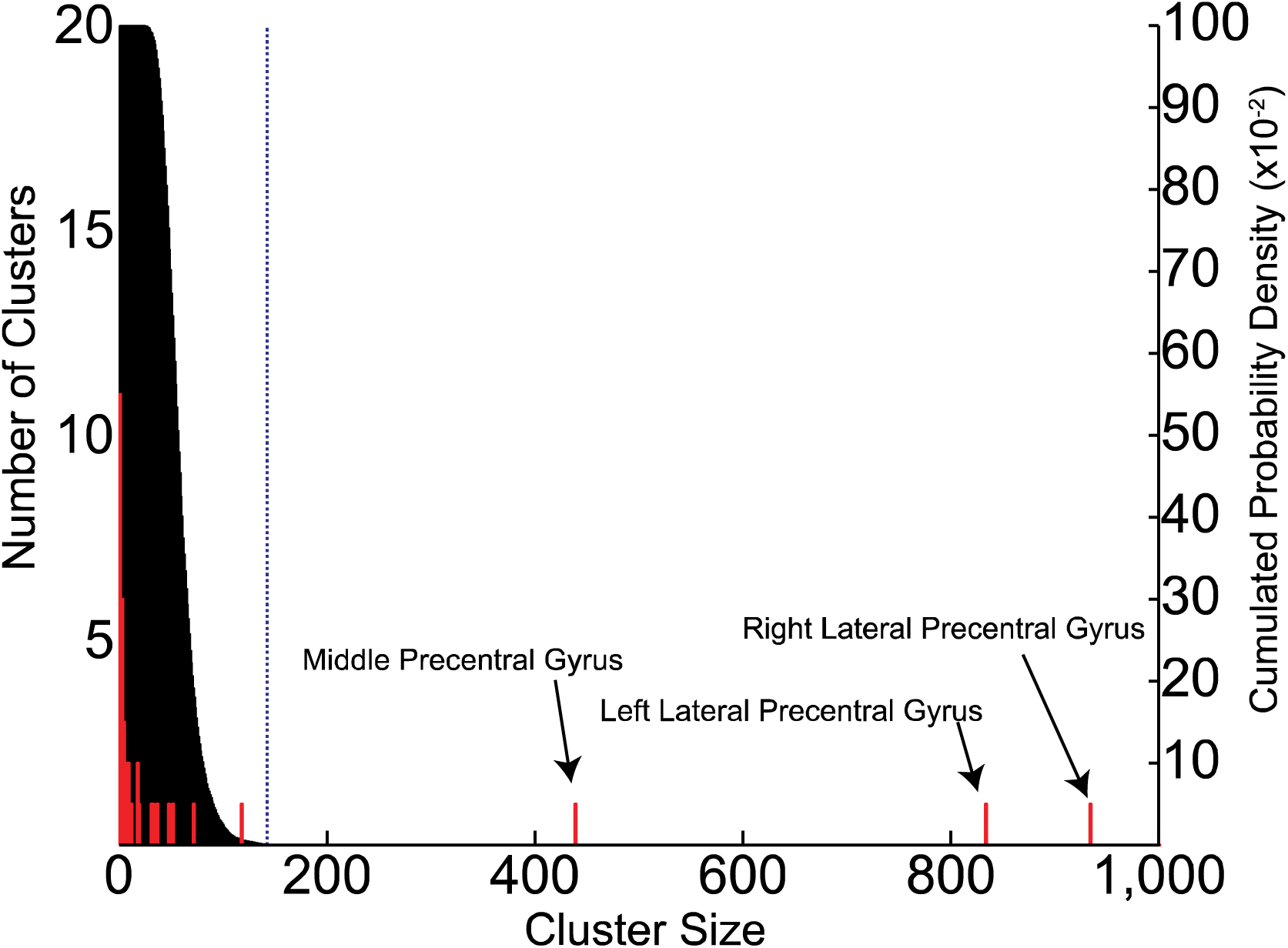
Cumulated probability density of maximum cluster size under the null hypothesis, *“The selected voxels were located randomly in the brain”* (black bars) and histogram of the sizes of the true relevant clusters (red bars). The vertical dotted line indicates the upper 0.1% (5% / 48) point of the null distribution (cluster size = 142). Three above-threshold clusters were identified, in the middle precentral gyrus (SMA; cluster size = 439), the lateral part of the precentral gyri in the right hemisphere (right M1/PMd; 934 voxels), and the lateral part of the precentral gyri in the left hemisphere (left M1/PMd; 834 voxels).

## Appendix B: Skull decoding

To verify that the success of decoding was not influenced by noise, we performed the decoding analysis with voxels outside the brain.

### Method

For each participant, the ROI for this analysis (Skull ROI) was first determined in MNI standard coordinates using the normalized T1-weighted anatomical image. The ROI included the voxels satisfying the following three criteria: 1) the fMRI signal in the T1 anatomical image exceeded 100; 2) the z coordinate was zero or positive; and 3) the distance from the nearest brain voxel was more than 8 mm. This ROI was then converted to the individual brain coordinates.

These criteria were used to ensure that the ROI did not include brain regions when converted to the individual brain coordinates, but included voxels in the head structure that could have been sufficient to identify signal changes caused by confounding noise, such as head movement artifacts. The number of voxels used in this analysis was 7,261 ± 591.

### Results

The decoding accuracy (Appendix Figure B.1) was near chance, except for Volumes #5 (Execution Period) and #6 (first volume in ITI). The t-test identified significantly higher decoding accuracies (*p*_*8*_ < 0.05 with two-tailed t-test with Bonferroni correction for multiple comparisons) for Volumes #5 (t_11_ = 7.08, *p*_*8*_ = 0.0002) and #6 (t_11_ = 5.45, *p*_*8*_ = 0.0016), but not for the other volumes (t_11_ < 2.9, *p*_*8*_ > 0.11).

### Discussion

Successful prediction based on out-of-the-brain voxels (Skull ROI) may suggest that Volumes #5 and #6 contained artifact signal irrelevant to the neural activity but informative about the selected hand. This was most likely due to artefactual signal changes caused by finger movement, because the response (button press) was performed during Volumes #5 and #6 in almost all trials.

Another possible explanation of the successful decoding from out-of-the-brain voxels is that the brain signals somehow leaked into the skull, such as phenomena such as N/2 ghosts, and the diffused brain signals led to the success of the prediction. However, this was probably not the main cause of the successful decoding for Volumes #5 and #6, because the decoding accuracy was near chance in Volumes #7 and #8, where the decoding accuracy was almost 100% in the whole-brain decoding (Figure 2A).

Thus, we argue that the skull decoding could successfully identify artefactual signal changes caused by finger movement, and importantly, Volume #4 was most likely free of such artifacts, since the mean decoding accuracy for Volume #4 in skull decoding was approximately at chance (50.8 ± 6.4%; t_11_ = 0.44, *p*_*8*_ = 5.3).

**Appendix Figure B.1:**
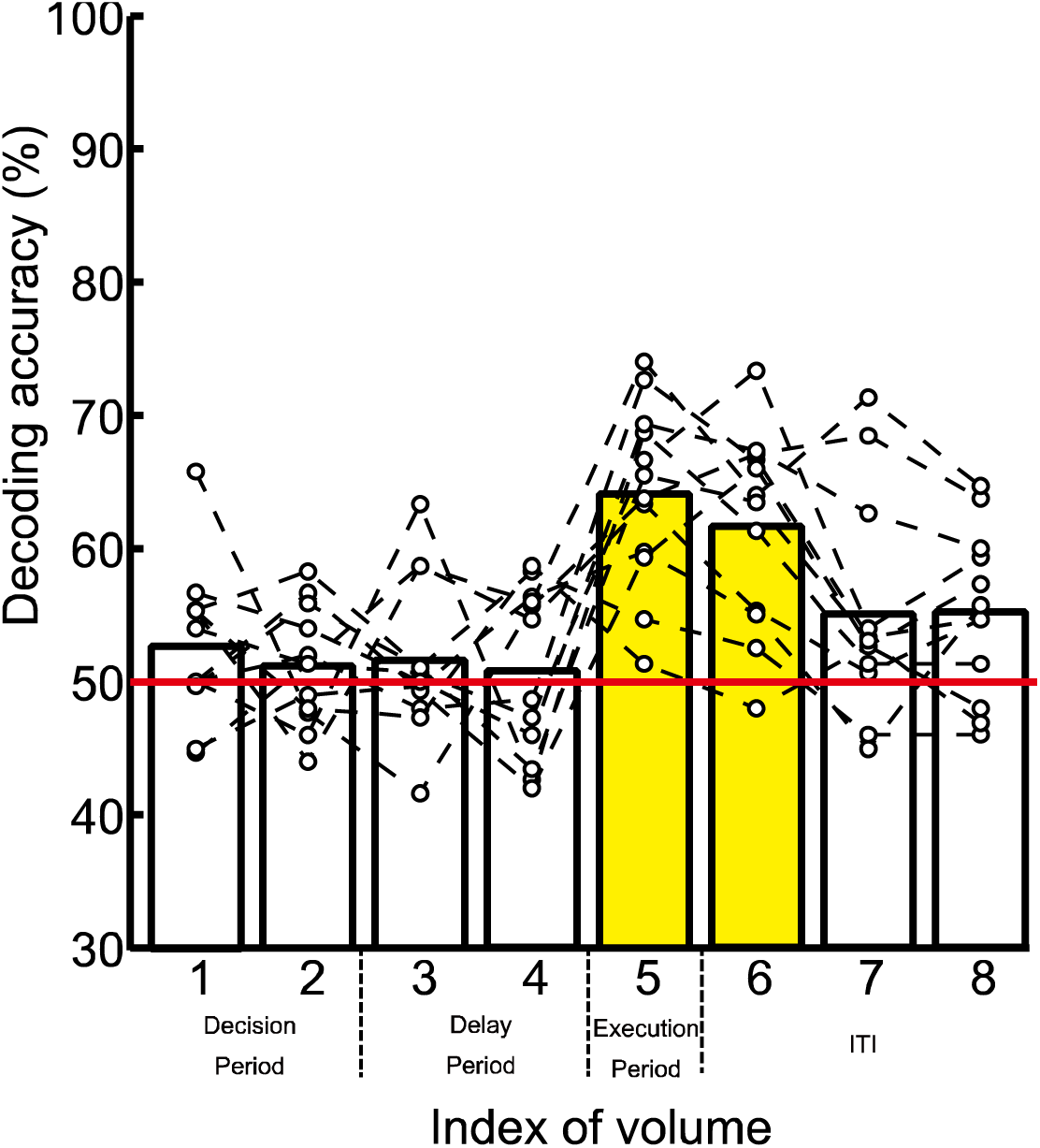
Decoding accuracies when voxels outside the brain were used (skull decoding). Bars indicate the mean decoding accuracy across participants, while white dots connected by dashed lines represent results from each participant. Yellow bars indicate the volumes for which t-tests revealed significantly higher-than-chance mean decoding accuracy (50%; red line), while white bars indicate a lack of significance.

## Acknowledgments

We would like to thank Dr. Nobuhiro Hagura, Center for Information and Neural Networks (CiNet), National Institute of Information and Communications Technology (NICT), for helpful suggestions on earlier versions of the manuscript. This work was supported by a Grant-in-Aid for Challenging Exploratory Research (JSPS KAKENHI 26560303) to author IN, Scientific Research on Innovative Areas “Embodied-brain” (JSPS KAKENHI No. 26120003) to EN, Grant-in-Aid for Specially Promoted Research (No. 24000012) to EN and Grant-in-Aid for Young Scientists (B) (JSPS KAKENHI 16K16649) to SH.

## Supplementary Material 1: Behavioral data analysis

We analyzed two aspects of behavioral data (sequence of the hand selection) to verify that the successful decoding is not due to biases in the participants’ hand selection.

### The balance of choices

A concern is that a trivial decoder can predict accurately if participants select one hand more frequently. For example, if a participant often selects his or her right hand, the decoding accuracy of a decoder that always predicts right hand movement (One-Selection Decoder) will be above 50%. However, we found that this was not the case in our experiment.

The mean probability that each participant selected the hand that was most frequently selected was 0.53 ± 0.02. Thus, the mean decoding accuracy of a One-Selection Decoder would have been 53%. We performed a paired permutation test with a procedure similar to that described in Appendix A.4, and found that the decoding accuracies obtained from the whole-brain decoding for Volume #4 and from the M1/PMd and SMA decoding were significantly better than those for the One-Selection Decoder (p = 0.0015).

Moreover, to confirm that the classifier performed equally well on the left and right hands, we performed a *χ*^2^ test of independence, and tested whether the difference in the correctness rates for the left and right hands was significant. The test did not identify a significant difference in any participant (*p* > 0.27, *χ*^2^ < 1.20).

Thus, we can rebut the suggestion that the accurate predictions reported in the text is due to bias of the participants’ hand selection.

### Carry-over of information from the previous trial

Another concern is that our successful prediction could have depended on the brain activation for the previous trial, rather than preparation for the current trial. If the hand selection could be predicted by observing selection of the previous trial, the brain activation observed in the previous trial would be useful for prediction of the current trial. Psychologists have shown that humans tend to choose candidates alternatively when they are instructed to choose one of two candidates randomly (e.g., Gerjuoy and Gerjuoy, 1964). In fact, in our study participants tended to alternate hands: they selected the opposite hand in 66.4% ± 3.9% of trials. Thus, if the decoder perfectly predicted the effector used in the previous trial, it could predict the effector in the upcoming trial in 66.4% ± 3.9% of the trials. We refer to this the Opposite Predictor.

However, for the following reasons, we are of the opinion that the successful decoding was not a result of contamination by brain activation from the previous trial. First, Volume #4 was acquired 12 s after the end of the previous trial. Thus, the effect of the hemodynamic response for the previous trial should have been negligible (Friston et al., 1994). Second, the decoding accuracies obtained from the whole-brain decoding for Volumes #1 and #2 were at almost chance levels, which suggests that the fMRI signal representing brain activation for the previous trial did not contribute to the prediction even at the beginning of the trial. Finally, there was no correlation between the decoding accuracy of the Opposite Predictor and the decoding accuracy obtained from the whole-brain for Volume #4 (R = 0.044, test of no correlation *p* = 0.89).

## Supplementary Material 2: Spatial distribution of representation of each hand

### Method

To identify the spatial distribution of the neuronal representation of each hand in M1/PMd and SMA, we again analyzed the spatial distribution of the voxel weights in the M1/PMd and SMA ROIs obtained in the whole-brain decoding.

First, for each participant, we identified the voxels that were assigned with positive weight values in more than half of the 10 validation tests (positively-weighted voxels). As in the procedures described in section 2.7, the locations of these voxels were transformed into MNI coordinates and smoothed. The voxels identified by this procedure were defined as “right hand voxels,” because higher values of positively-weighted voxels push the decoder toward a label prediction of 1 (right hand; Hirose et al., 2015). The voxels that were assigned with negative weight values were also analyzed with the same procedures to identify the “left hand voxels.” Using t-tests, we compared the number of right hand voxels with left hand voxels in the right and left M1/PMd relevant clusters.

For the SMA ROI, we separated the ROI into right (x > 0 in MNI coordinates) and left (x < 0) hemispheres and performed the same analysis. Finally, for visualization, we identified the voxels which were left/right hand voxels in more than 2 participants (Supplementary Figure A).

### Results

We observed that hand representations were clearly localized to the contralateral hemisphere in both M1/PMd and SMA (Supplementary Figure); i.e., the majority of the left hand voxels were located in the right hemisphere (right M1/PMd; 850 ± 499; left M1/PMd; 112 ± 126; right SMA; 190 ± 145; left SMA; 14 ± 16), while the majority of the right hand voxels were located in the left hemisphere (right M1/PMd; 105 ± 179; left M1/PMd; 847 ± 469; right SMA; 17 ± 26; left SMA; 196 ± 136). We also identified a significant difference between right and left hand voxels in both the M1/PMd and SMA ROIs of both hemispheres (t_11_ > 4.3; *p*_*4*_ < 0.005).

**Supplementary Figure:**
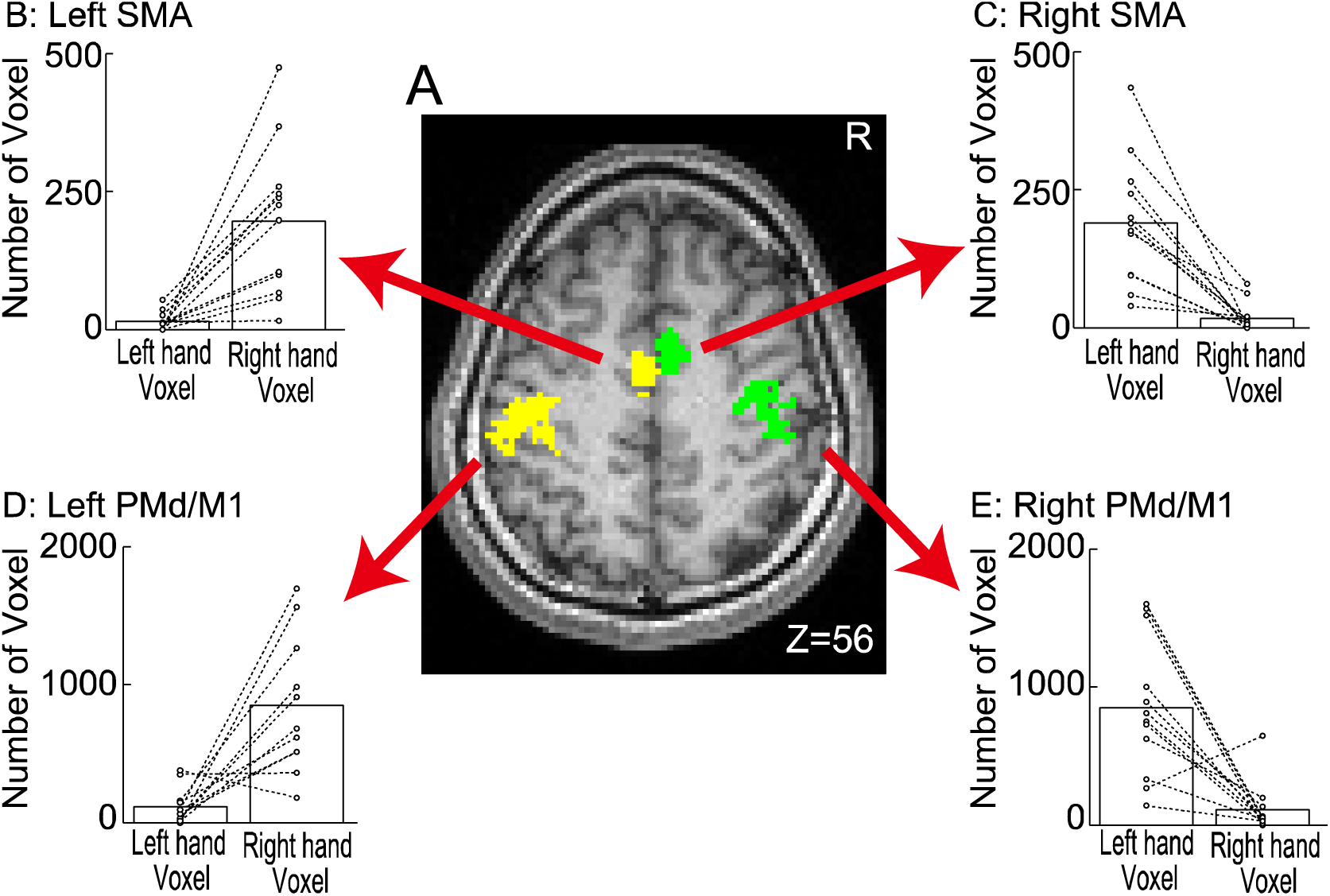
Right and left hand voxel distribution. A) Each hand’s representation in the right and left hemispheres of the relevant clusters. Green and yellow sections represent voxels that were left and right hand voxels in more than 2 participants, respectively. B-E) Number of left and right hand voxels for each region (B: Left SMA, C: Right SMA, D: Left M1/PMd, E: Right M1/PMd). Bars indicate mean numbers across participants. Dots connected with a dashed line indicate results from each participant.

**Supplementary Table 1:**
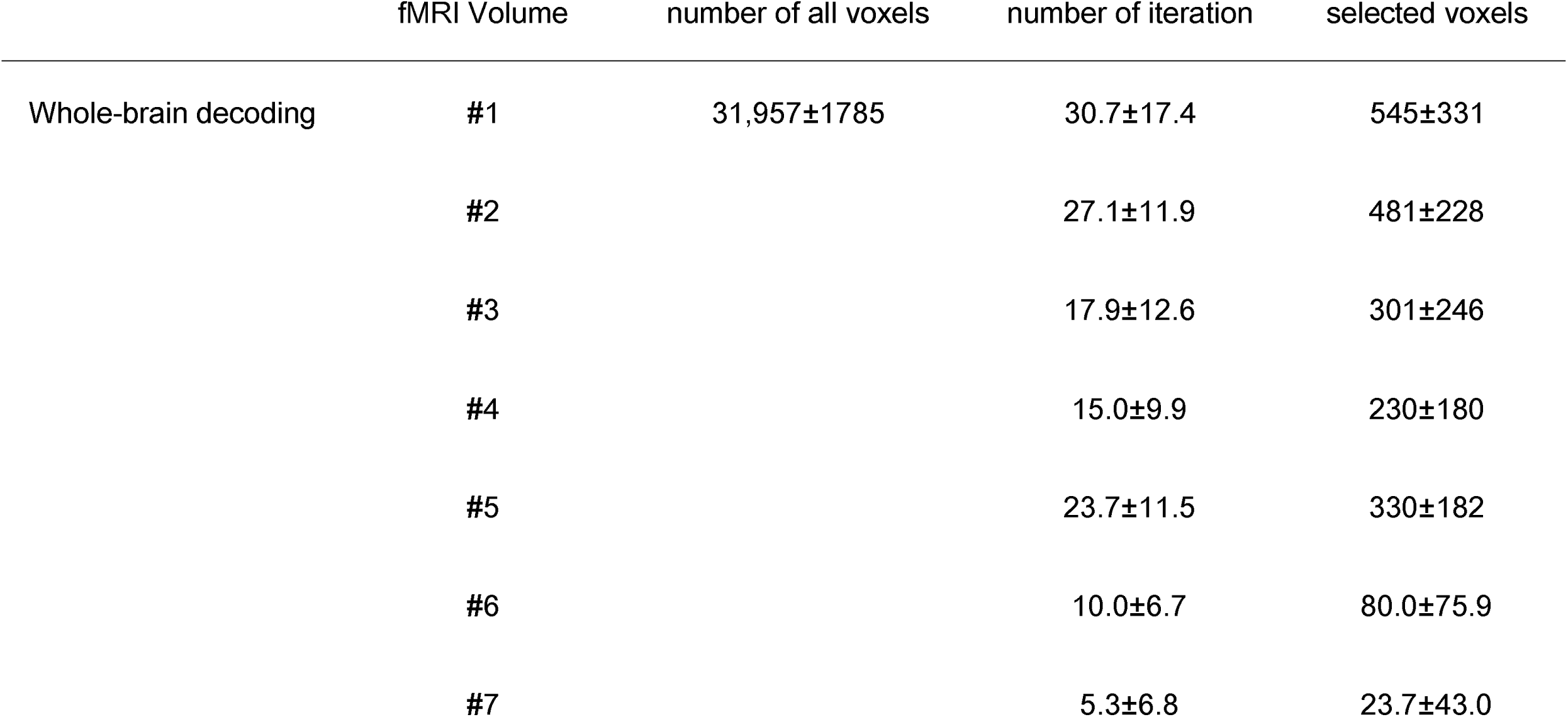
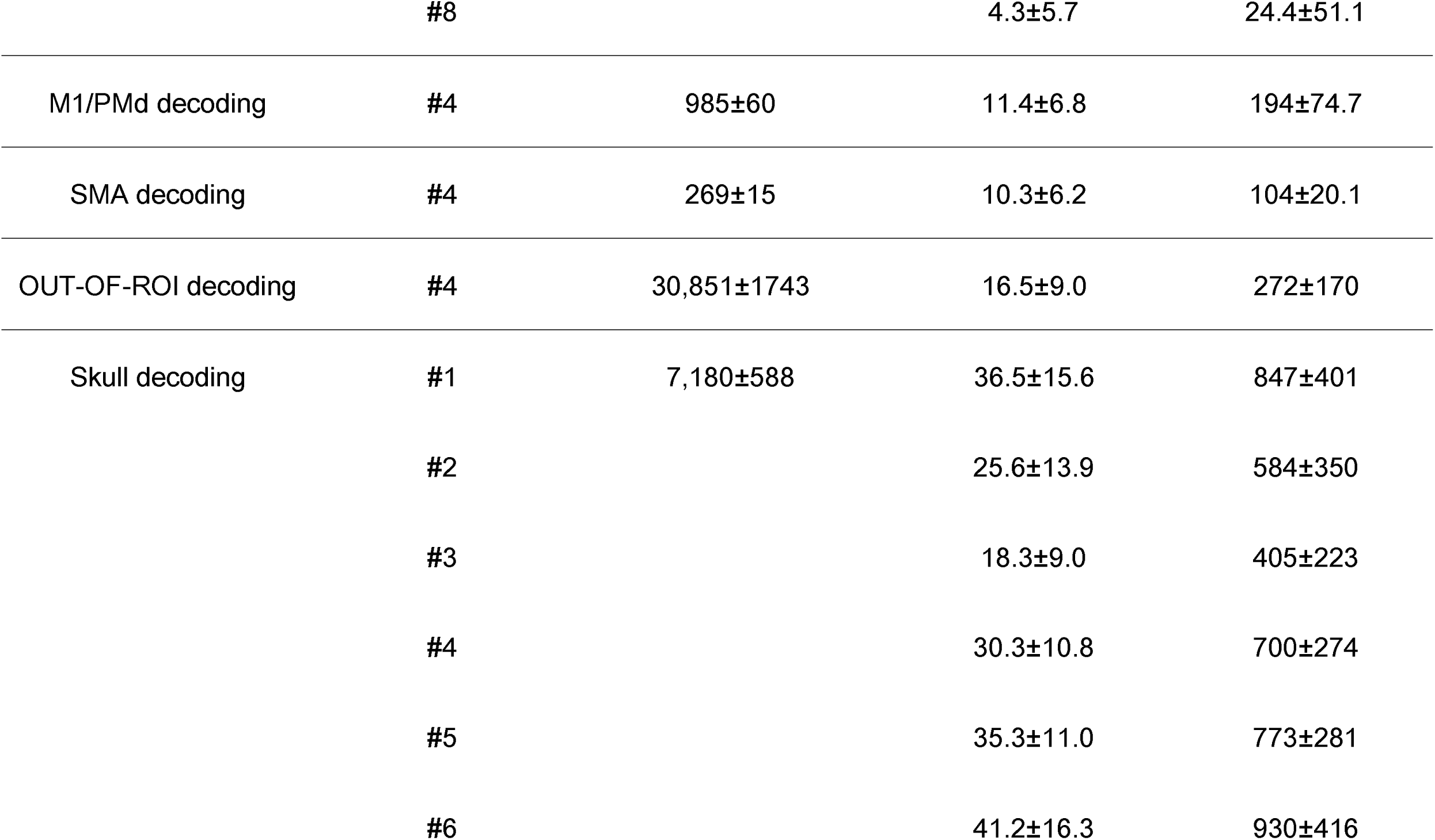
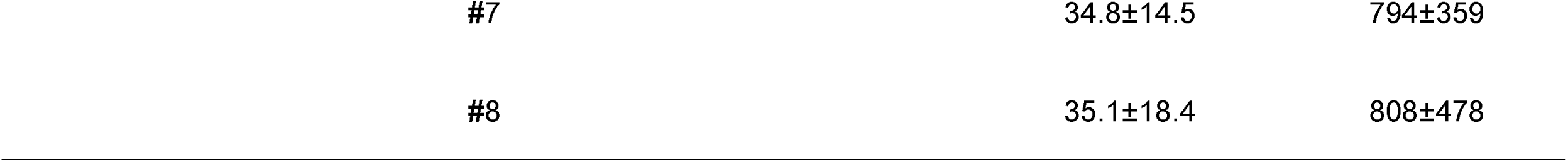
Mean number of iSLR iterations and selected voxels for each decoding analysis.

